# Powerful and interpretable control of false discoveries in differential expression studies

**DOI:** 10.1101/2022.03.08.483449

**Authors:** Nicolas Enjalbert-Courrech, Pierre Neuvial

## Abstract

**Motivation:** The standard approach for statistical inference in differential expression (DE) analyses is to control the False Discovery Rate (FDR). However, controlling the FDR does not in fact imply that the proportion of false discoveries is upper bounded. Moreover, no statistical guarantee can be given on subsets of genes selected by FDR thresholding. These known limitations are overcome by post hoc inference, which provides guarantees of the number of proportion of false discoveries among arbitrary gene selections. However, post hoc inference methods are not yet widely used for DE studies.

**Results:** In this paper, we demonstrate the relevance and illustrate the performance of adaptive interpolation-based post hoc methods for DE studies. First, we formalize the use of permutation-based methods to obtain sharp confidence bounds that are adaptive to the dependence between genes. Then, we introduce a generic linear time algorithm for computing post hoc bounds, making these bounds applicable to large-scale DE studies. The use of the resulting Adaptive Simes bound is illustrated on a RNA sequencing study. Comprehensive numerical experiments based on real microarray and RNA sequencing data demonstrate the statistical performance of the method.

**Availability:** A cross-platform open source implementation within the R package `sanssouci` is available at https://pneuvial.github.io/sanssouci/.

## 1. Introduction

Two-sample comparison problems are ubiguitous in genomics. The most classical example is the case of differential expression studies, where the goal is to pinpoint genes (or transcripts) whose average expression level differ significantly between two known populations, based on a sample of expression measurements from individuals from these populations. A classical strategy to identify *differentially expressed* (“DE”) genes is to test, for each gene, the null hypothesis that its average expression is identical in both populations. DE genes are then defined as those passing some significance threshold, after accounting for the fact that many tests are performed simultaneously.

The state of the art approach to large-scale multiple testing is to control the False Discovery Rate (FDR). Introduced by Benjamini and Hochberg (1995), the FDR is the expected proportion of wrongly selected genes (false positives) among all selected genes. The most widely used method to control FDR is the Benjamini-Hochberg (BH) procedure, which has been shown to control FDR when the hypotheses corresponding to the non-differentially expressed genes are independent or satisfy a specific type of positive dependence called PRDS (Benjamini and Yekutieli, 2001). PRDS is widely accepted as a reasonable assumption in differential gene expression (DGE) studies and in genomic studies in general, see e.g. Goeman and Solari (2014). However, there exist two major caveats to the practical use and interpretation of FDR in genomics. Let us assume that we have obtained a list *R* of genes called DE by a FDR-controlling procedure applied at level *q.*

### Practical use: FDR of gene subsets is not controlled

As noted by Goeman and Solari (2011), the statement FDR(R) ≤ q applies to the list *R* only, and no further statistically valid inference can generally be made on other gene lists. However, a common practice is to manually curate this list by adding or subtracting genes, based on some external or priori knowledge (such as the knowledge of gene sets or pathways). A typical example is the case of volcano plots (Cui and Churchill, 2003), where one selects those genes passing both a significance threshold and a threshold on the fold change (difference of average gene expression on the log scale), see Figure 3. Ebrahimpoor and Goeman (2021) have recently shown in an extensive simulation study that this type of double filtering strategy yields inflated false discovery rates.

### Interpretation: FDR control is not FDP control

The statement FDR(*R*) ≤ *q* is often misinterpreted as “the proportion of false discoveries (FDP) in *R* is less than *q*”. In fact, the FDP is a *random* quantity, and FDR(*R*) ≤ *q* only implies that the *average FDP over hypothetical replications* of the same genomic experiment and *p*-value thresholding procedure, is upper bounded by *q*. This distinction would not matter much if gene expressions were statistically independent: indeed, as the number *m* of tests tend to infinity, the FDP concentrates to the corresponding FDR with a typical parametric convergence rate: *m*^-1/2^ (Neuvial, 2008). Unfortunately, as the dependence increases, the FDP distribution becomes strongly asymmetric and heavy-tailed, as illustrated by the numerical experiments reported in Neuvial (2020, Fig. 2.1).

The notion of *post hoc inference* has been introduced by Goeman and Solari (2011) to address these limitations. Building on earlier works by Genovese and Wasserman (2006), Goeman and Solari (2011) have obtained confidence bounds for the FDP in *arbitrary, multiple and possibly data-driven subsets of hypotheses* using the theory of closed testing (Marcus *et al.*, 1976). In practice, *Simes post hoc bounds* are recommended in Goeman *et al.* (2019), as they are valid under the PRDS assumption and can be calculated efficiently. Simes post hoc bounds have recently been popularized in genomics by Ebrahimpoor and Goeman (2021), but also in neuroimaging studies by Rosenblatt *et al.* (2018), where this approach has been called “All-resolutions inference” (ARI).

Despite their very attractive theoretical properties, post hoc methods are not yet widely known and used for addressing multiple testing situations in genomics, where controlling FDR via the BH procedure remains standard. Two possible reasons for this situation are that contrary to the BH procedure for FDR control, the Simes post hoc bound for post hoc inference is (i) typically conservative in genomic applications, and (ii) its construction based on closed testing may be difficult to understand for practitioners.

An alternative construction of post hoc bounds that has been proposed in Blanchard *et al.* (2020) and further explored in Blanchard *et al.* (2021); Durand *et al.* (2020). This strategy can yield sharper bounds, by an adaptation to the statistical dependency between tests using permutations, and to the sparsity of the signal using a step-down principle. The goal of the present paper is to popularize the use of the post hoc bounds introduced in Blanchard *et al.* (2020) in the context of DE studies. Besides providing a self-contained introduction to interpolation-based post hoc inference (Section 2), the main contributions of this paper can be summarized as follows:

1. Powerful post hoc bounds can be obtained by leveraging permutation methods in order to adapt to the unknown dependency structure of a given data set (Section 3);
2. Generic interpolation-based post hoc bounds can be computed in linear time (Section 4);
3. Application of the resulting “adaptive Simes” method to a specific RNA seq DE study yields more interpretable results than those derived from FDR control, and sharper bounds than Simes post hoc bounds (Section 5);
4. Comprehensive numerical experiments based on real genomic data demonstrate the statistical performance of the method (control of the target risk, and statistical power) for DE studies, both for microarray and sequencing data sets (Section 6).

These developments are implemented in the R package `sanssouci` available from https://pneuvial.github.io/sanssouci/. The code used for the numerical experiments s available from the package source.

## 2. Interpolation-based post hoc inference

We consider a DE study with *m* features. These features are called genes for simplicity, but the methods described below are also applicable more generally. For now, we only assume that a *p*-value is available to test the differential expression of each gene. The vector of *p*-values is denoted by (*p*_1_,…,*p_m_*). More specfic assumptions on how these *p*-values are obtained are given in Section 3.

### 2.1. Objective: post hoc bounds

For a given subset *S* of genes called DE, we denote by FP(*S*) the number of false positives in S, that is, the number of genes in *S* that are not truly DE. Our goal is to find a function 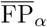 such that with high probability, 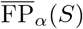 is larger than the number of false positives in *S*:

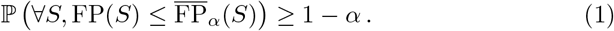

Following Goeman and Solari (2011), a function 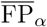 satisfying (1) will be called an *α*-level *post hoc upper bound on the number of false positives.* Post hoc inference can be equivalently formulated in terms of upper bounds on the 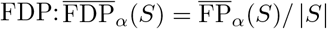, or in terms of lower bounds on the number or proportion of true positives: 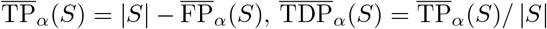.

### 2.2. Strategy: JER control and interpolation

The bounds studied in this paper rely on a multiple testing risk called the Joint Error Rate (JER) and introduced in Blanchard *et al.* (2020). Given a non-decreasing family of thresholds **t** = (*t_k_*)_*k*=1…*K*_,

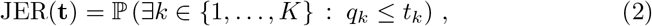

where for *k* = 1,…,*K*, *q_k_* denotes the *k*-th smallest *p*-value among the set of truly non-DE genes (that is, true null hypotheses). A key result is that any family **t** such that JER(**t**) ≤ *α* yields an associated *α*-level post hoc bound, by the following simple interpolation argument.

#### Proposition 1

(Interpolation-based post hoc bound (Blanchard *et al.*, 2020), Proposition 2.3). *If* **t** = (*t_k_*)_1≤*k*≤*K*_ *controls JER at level α, then (1) is satisfied for the bound*

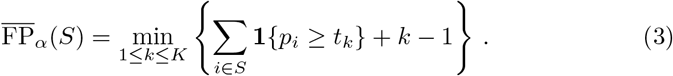

A proof of Proposition 1 is given in Appendix B for completeness.

### 2.3. Simes post hoc bounds

An important example is the Simes family **t**^S^(*α*), defined by 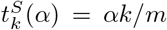 for all *k*. The Simes (1986) inequality ensures that JER(**t**^S^(*α*)) ≤ *α* as soon as the *p*-value family is PRDS (Sarkar *et al.*, 2008). As noted by Blanchard *et al.* (2020), the post hoc bound then derived by Proposition 1 coincides with the Simes post hoc bound introduced in Goeman and Solari (2011).

Although the Simes inequality is sharp when the *p*-values are independent, it is increasingly conservative as the dependence between tests gets stronger (Blanchard *et al.*, 2020, Table 1). The associated JER control and post hoc bound naturally inherit this conservativeness (as illustrated in the numerical experiments of Sections 5 and 6). In order to address this conservativeness issue, it is useful to note that for λ > 0, the JER of the Simes family **t**^S^(λ) can be written as

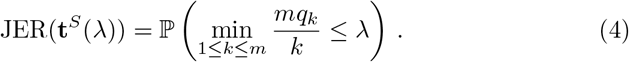

**Table 1.** *Post hoc bounds on BLCA data set for ARI and the proposed permutation method, for gene selections S illustrated in Figures 2 (target FDP*= 0.1) *and 3 (genes filtered by *p*-value and fold-change).*

|  | Confidence curve (Fig. 2) |  | Volcano plot (Fig. 3) |  |
| --- | --- | --- | --- | --- |
|  | ARI | Adaptive Simes | ARI | Adaptive Simes |
| Number of genes in $S$ | 781 | 1064 | 569 | 569 |
| $\overline{\text{TP}}_\alpha(S)$ | 703 | 958 | 456 | 492 |
| $\text{FDP}_\alpha(S)$ | 0.1 | 0.1 | 0.199 | 0.135 |

In view of (4), a natural idea in order to obtain a tight JER control is to select the largest λ such that JER(**t**^*S*^(λ)) ≤ *α*. This idea is the basis of the calibration method described in Section 3.

## 3. JER calibration by permutation

The JER defined in (2) only depends on the joint *p*-value distribution of true null hypotheses. Although this distribution is unknown in practice, in two-group DGE studies, it can be approximated by permuting the group labels. Accordingly, the first step of our calibration method is to build a *B* × *m* matrix *P* of permutation *p*-values: *P_bi_* is the *p*-value of the test of gene *i* associated to the *b*-th permutation of the group labels. This is illustrated in the first panel of Figure 1.

**Fig 1.**
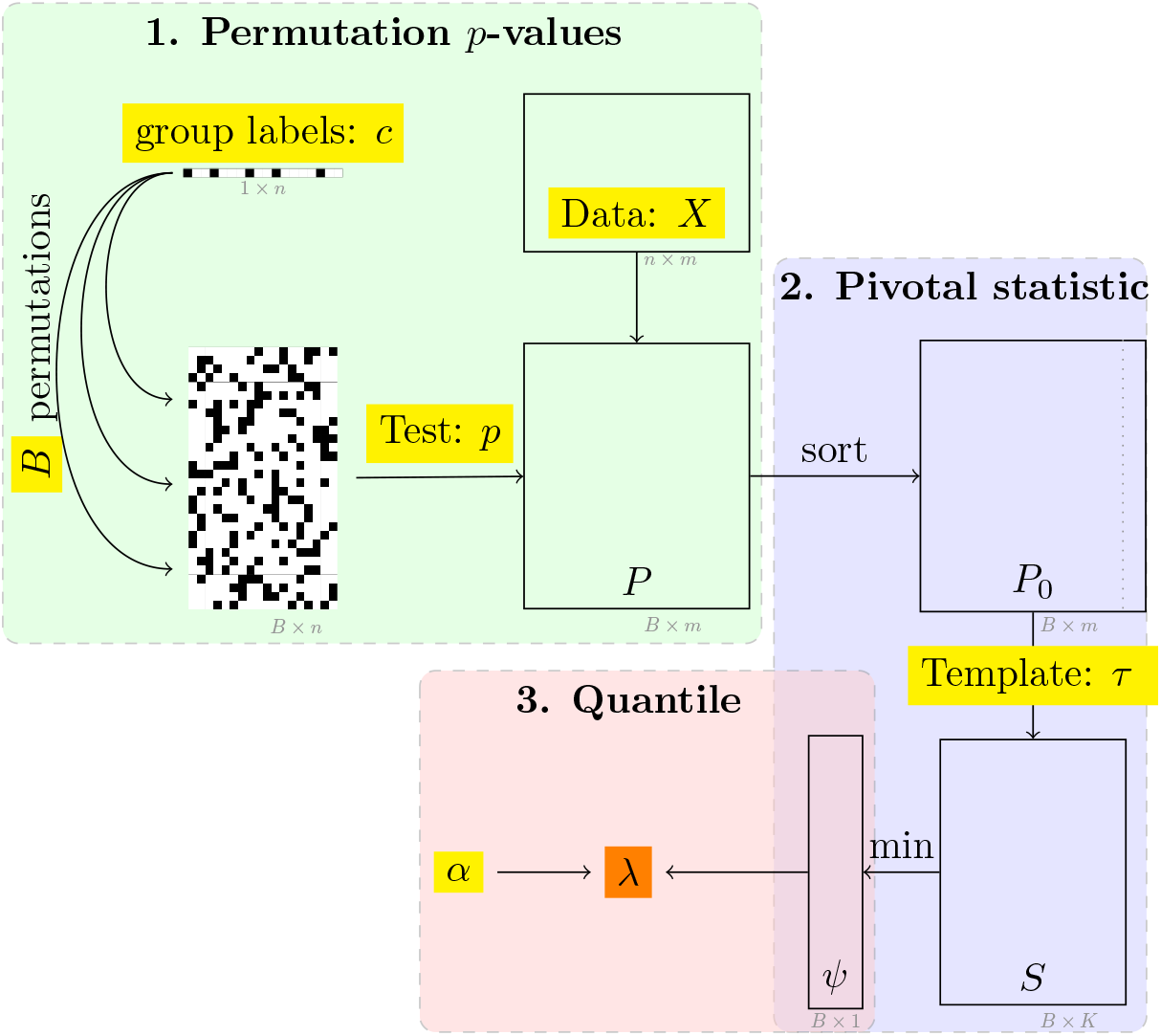
Illustration of the three main steps of permutation-based JER calibration. The output is highlighted in orange, and inputs are highlighted in yellow. The input data is in the form of a n × m gene expression matrix X and a binary vector c of n group labels, specifying which observations belong to each of the two populations to be compared. The parameters are the target JER level α and the number B of permutations, the p-value function to perform the test and the template τ.

The next steps of the calibration are best explained in the particular case of the Simes family. Indeed, by (4), JER(**t**^*S*^(λ)) is the value of the cumulative distribution function of *ψ* = min_1≤*k*≤*m*_ *mq_k_*/*k* at λ. Accordingly, the calibration method proceeds by calculating *B* samples from the “pivotal statistic” *ψ*, and the output is the quantile of order α of these statistics.

The method as described in Figure 1 covers not only the case of the Simes family, but any family *τ*(λ) = (*τ_k_*(λ))_*k*=1…*K*_ where the *τ_k_* are invertible functions. Following Blanchard *et al.* (2020, 2021), such a family is called a *template*. A more formal description of this calibration algorithm is given in Algorithm 2. Note that for simplicity, we have described here a “single-step” version of the calibration algorithm. We have also implemented a “step-down” version: it is a slightly more powerful algorithm that is also adaptive to the unknown proportion of true null hypotheses (Blanchard *et al.*, 2020, Proposition 4.5).

### Validity

Theorem 1 in Blanchard *et al.* (2021) ensures that this calibration method yields JER(λ) ≤ *α*, for tests whose *p*-value for a given gene depends on the data only via its own expression values. In particular, this is the case for two-sample Student tests or Wilcoxon rank sum tests, which can be used for microarray and RNA sequencing (RNAseq) DE studies, respectively. The theory developed in Blanchard *et al.* (2021) is also valid for one-sample tests: in this case, permutation *p*-values at step 1 are replaced by sign-flipping *p*-values.

### Complexity

Assuming a linear time complexity *O*(*n*) to perform the test of one single null hypothesis, the overall time complexity of the calibration method is *O*(*mB*(*n* + log(*m*))). Indeed, the most costly step is the calculation of *P*_0_, which involves mB tests followed by *B* sorting operations on a vector of size *m*. The overall space complexity is *O*(*m*(*B* + *n*)).

Figure 1 also illustrates the modularity of Algorithm 2, where the three main steps are highlighted in different colors. This modularity is important in practice. For example, it makes it possible to obtain the result for several values of α without re-computing the permutation matrix *P*_0_. This modularity is also useful for the computational efficiency of the above-mentioned step-down version of the calibration algorithm.

## 4. Linear time interpolation-based post hoc bound

Post hoc bounds can be used for multiple gene selections *S* without compromising the corresponding error control. For post hoc inference to be applicable in practice 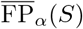 must be computed efficiently.

A naive implementation of the bound 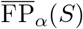 defined in (3) would require *s*^2^ operations (where *s* is the size of *S*) by performing a loop on both *k* = 1,…, *s* and *i* ∈ *S* in order to calculate *v_k_* (*S*) ∑*i*∈*S* **1**{*p_i_* ≥ *t_k_*} + *k* – 1 for all *k*. This induces a quadratic worst case time complexity *O*(*m*^2^), which is achieved when evaluating 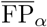 on the set of all genes. A quadratic time complexity for a single set is too slow for DE studies with *m* ≥ 10, 000. Moreover, a useful application of post hoc bounds is to build the false positive confidence curve associated to *S*, that is, all the bounds 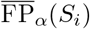 for *i* = 1… *s*, where *S_i_* is the index set of the i smallest *p*-values in *S*. Using the above naive algorithm, this would require *O*(*s*^3^) operations, implying a cubic worst case time complexity *O*(*m*^3^) to build the false positive confidence curve associated to all hypotheses.

In contrast, Algorithm 1 computes 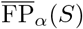 in linear time and space *O*(*s*) for a given *S* In fact, it even outputs the entire false positive confidence curve associated to SFor example, the largest set *S* such that 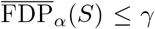 is then obtained in linear time and space for any user-defined *γ*. This complexity cannot be improved since the size of the output vector is *s*. The validity of algorithm 1 relies on the following formulation for 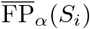.

**Algorithm 1.**
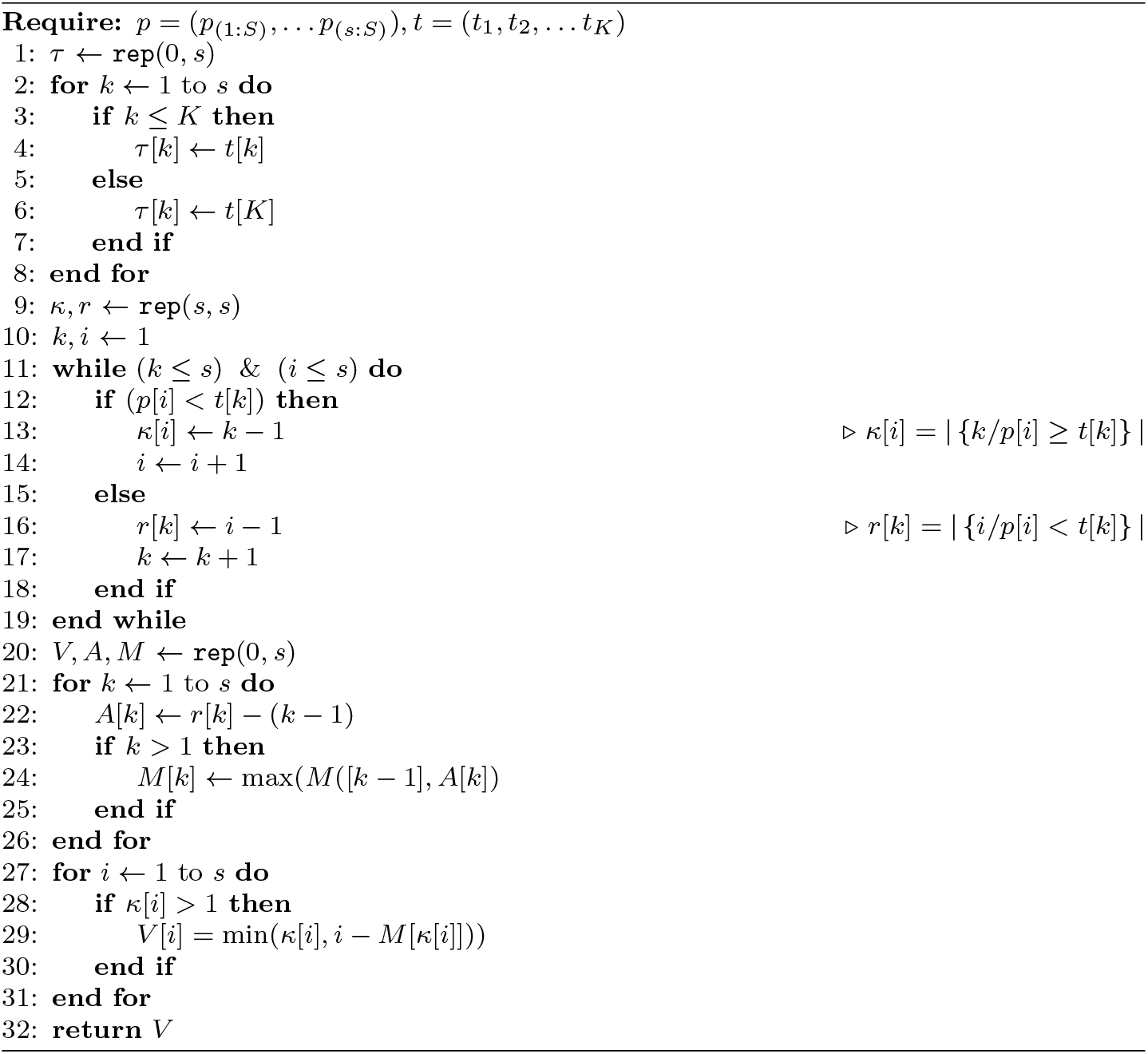
Linear algorithm for interpolation-based post hoc bounds.

### Proposition 2.

For *i* ∈ {1,…, *s*}, *let S_i_ be the index set of the i-th smallest p-value in S, and κ_i_* = ∑_*k*=1_…*s* **1**{*p_i_* ≥ *t*_*k*Λ*K*_}. *Then*

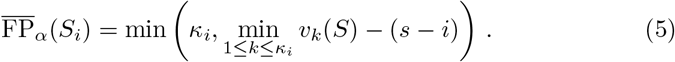

Proposition 2 is proved in Appendix C. The fact that 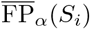 depends on *i* only via *κ_i_* but not *S_i_* in (5) is crucial for obtaining a linear time complexity. The properties of Algorithm 1 can be summarized as follows:

### Corollary 1

(Validity and complexity of Algorithm 1). *Algorithm* 1 *returns the vector* 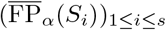 in *O*(*s*) *time and space complexity.*

*Proof of Corollary* 1. **Valididy.** The for loop at lines 1-8 stores the thresholds (*t*_*k*Λ*κ*_) for *k* ∈ {1,…,*s*}. The while loop at lines 11-19 outputs both (*κ_i_*)_*i*∈*S*_ and (*r_k_*)_1≤*k*≤*K*_, where *r_k_* = |**∑**_*i∈S*_ **1**{*p_i_* < *t_k_*}|. Noting that *r_k_* = *s* – *v_k_*(*S*) + (*k* − 1), the **for** loop at lines 21-26 outputs *M_k_* = max_*k*′≤*k*_ *s* – *v*_*k′*_(*S*), that is, *M_k_* = *s* – min_*k*′≤*k*_ *v_k′_*(*S*). Thus, the **for** loop at lines 27-31 outputs 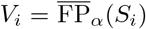 by Proposition 2.

### Complexity

All the vectors stored within the algorithm are of size *s*, so the space complexity of Algorithm 1 is *O*(*s*). For the time complexity, the (*κ_i_*)_*i*_ and (*r_k_*)_*k*_ are calculated within a single while loop of size *s*, in which exactly one of *i* or *k* is incremented at each step. The rest of the algorithm consists of two for loops of size *s* consisting of *O*(1) operations.

## 5. Urothelial Bladder Carcinoma data set

In this section, we focus on an Urothelial Bladder Carcinoma (BLCA) RNA sequencing data set from the Cancer Genome Atlas Research Network *et al.* (2014). This data set is available from the R/Bioconductor package `curatedTCGAData`. For convenience it has also been made available in the R package `sanssouci.data`. This data set consists of gene expression measurements for *n* = 270 patients, classified into two subgoups: stage II (*n*_0_ = 130) and stage III (*n*_1_ = 140). Bladder cancer stages range from 0 to IV, quantifying how much the cancer has spread. We have filtered out unexpressed genes, here defined as those for which raw expression counts were lower than 5 in at least 75% of the patients. This results in *m* = 12, 534 genes. To identify DE genes between the stage II and stage III populations, we test for each gene the null hypothesis that the gene expression distribution is identical in the two populations. The calibration method described in Section 3 is performed using a Wilcoxon (1945) rank sum test (also known as Mann and Whitney (1947) test) with the Simes template, with *B* = 1, 000 permutations and target risk (JER) set to *α* = 10%. The resulting method is called the **Adaptive Simes** method.

### 5.1. Confidence curves

In the absence of prior information on genes, a natural idea is to rank them by decreasing statistical significance. Post hoc methods provide lower confidence curves on the number (or proportion) of true positives (truly DE genes) among the most significant genes. Such curves are displayed in Figure 2 for the BLCA data set. The black lines in Figure 2 are 1 – *α* = 90% confidence curves obtained by the Adaptive Simes method. Upper bounds on FDP and lower bounds on FP are displayed in the left and right panels, respectively. For reference, the corresponding curves obtained by ARI are displayed in gray; they are almost identical to the ones obtained from the original bound of Goeman and Solari (2011) (Equation (3)), see Section D.

**Fig 2.**
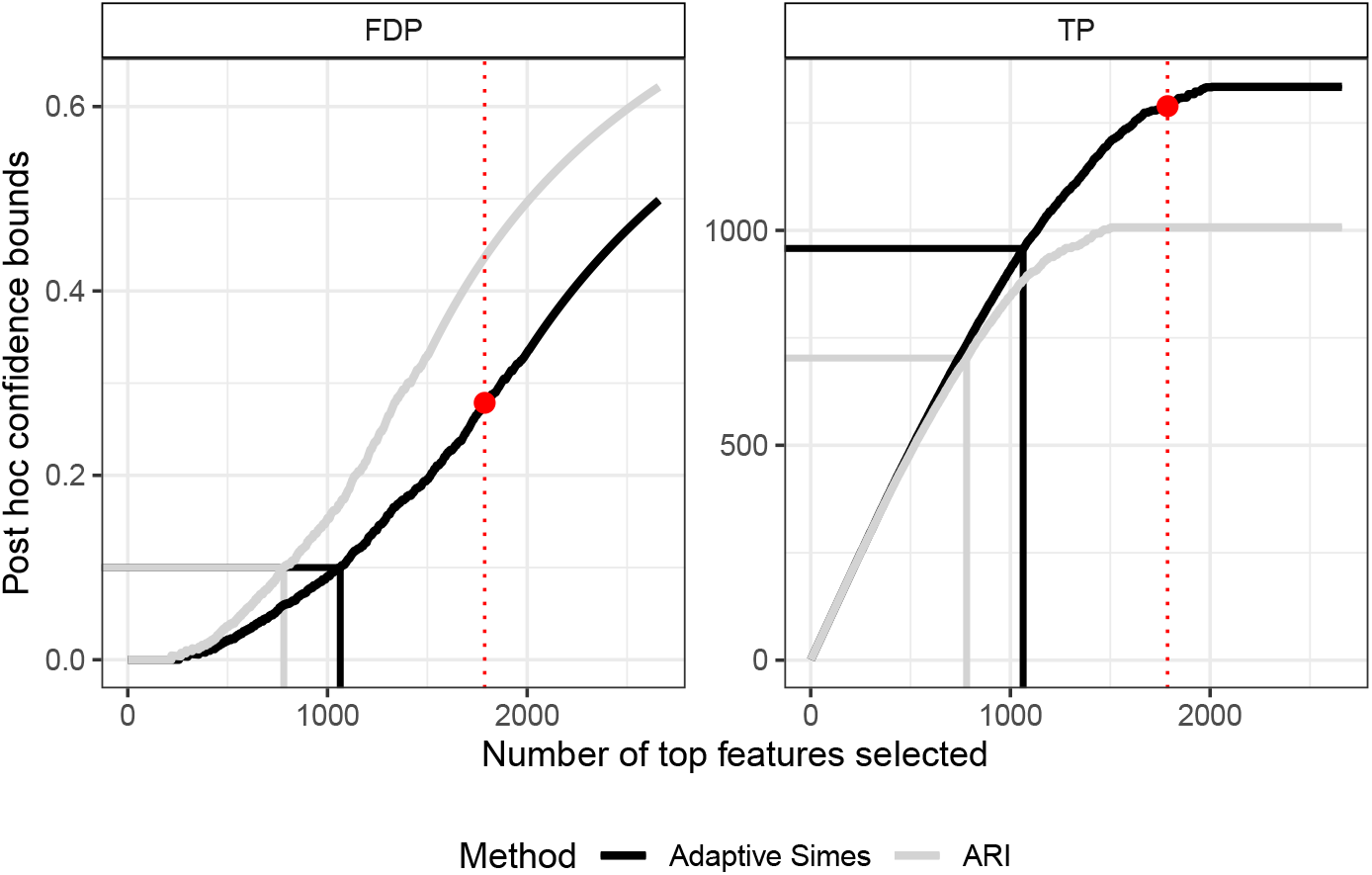
*90% confidence curves on “top k” lists for the Urothelial bladder carcinoma data set. Left: upper bound on the False Discovery Proportion (FDP); right: lower bound on the number of true positives (TP). The adaptive Simes bound (black curves) outperforms ARI (gray curves). For reference, the set of* 1, 787 *genes called DE by the BH(0.05) procedure is represented by a red dot.*

**Fig 3.**
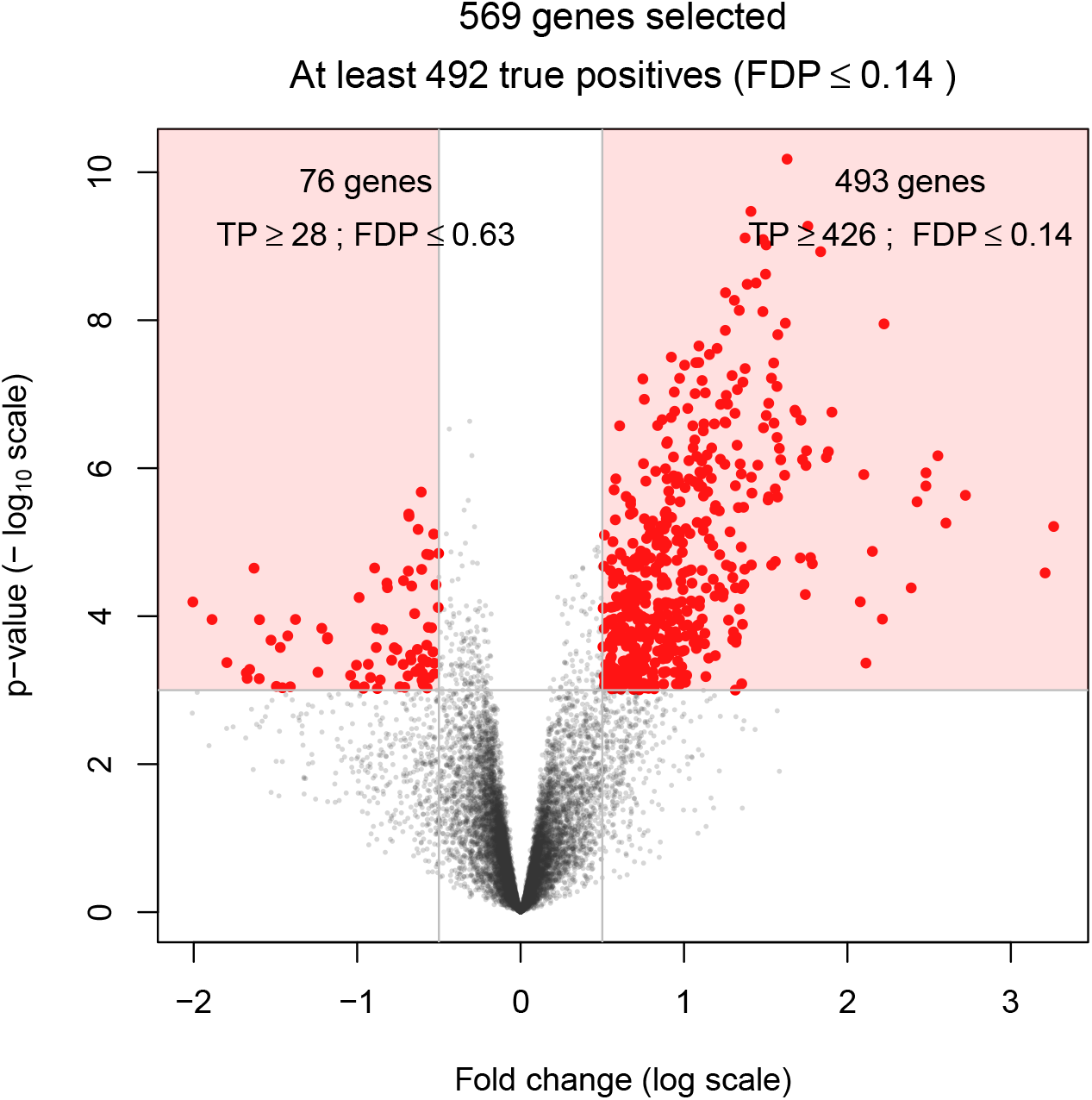
*Volcano plot for the urothelial bladder carcinoma data set. Each dot corresponds to a gene, represented by its fold change (x axis) and p-value (y axis) on the log scale. Fold changes and p-values were obtained by the limma-voom method (Ritchie* et al., *2015). The* 569 *genes with p-value less than* 10^-3^ *and fold change larger than* 0.5 *are highlighted. The Adaptive Simes method ensures that at least* 492 *of these genes are true positives.*

#### Post hoc guarantees

Post hoc inference makes it possible to define DE genes as the largest set of genes for which the FDP bound is less than a user-given value *q*. The arbitrary choice *q* = 0.1 is illustrated in Figure 2, corresponding to the horizontal line in the left panel. The black lines in Figure 2 correspond to the set *S* of 1064 genes for which the adaptive Simes method ensures that FDP(*S*) ≤ *q*. This corresponds to at least 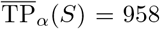 true positives (since 1 – 958/1064 = 0.1), as illustrated in the right panel.

#### Adaptation to dependence

The above example also illustrates the increase in power obtained by Adaptive Simes thanks to the calibration described in Section 3. Indeed, for an identical statistical guaranty (*FDP* ≤ 0.1), ARI yields a substantially smaller subset of 703 DE genes. More generally, the comparison between the black and gray curves in Figure 2 illustrates the gain in power obtained by using permutations methods to adapt to the dependence between genes.

#### Comparison to FDR control

For this data set, the BH procedure calls a set *S* of 1787 genes DE for a target FDR level of 0.05. As stated in Section 1, *the BH procedure does not provide guarantees on the FDP of these genes*, but only on their FDR, that is, the average FDP over hypothetical replications of the same genomic experiment and *p*-value thresholding procedure. In contrast, the Adaptive Simes bound guarantees (with 90% confidence) that the number of true positives in *S*^BH^ is at least 1, 289, or, equivalently, that the corresponding FDP is less than 0.279.

### 5.2. Volcano plots

Volcano plots are a commonly used graphical representation of the results of a differential expression analysis (Cui and Churchill, 2003), illustrated in Figure 3. Each gene is represented in two dimensions by estimates of its effect size (or “fold-change”, *x* axis) and significance (*y* axis). The fold change of a gene is generally defined as the difference between the average or median (log-scaled) gene expressions of the two compared groups. Its significance is quantified by – log_10_(*p*)-values for the test of its differential expression, where the “– log_10_”, transformation ensures that large values of y correspond to genes which are likely to be differentially expressed.

As noted by Ebrahimpoor and Goeman (2021), post hoc inference makes it possible to select genes of interest based on both fold change and significance, without compromising the validity of the corresponding bounds. Moreover, even if Wilcoxon tests have been performed for the *calibration of the post hoc bounds,* it is possible to rely on other statistics for the *selection genes of interest.* Figure 3 illustrates this idea by making a volcano plot based on the *p*-values and log-fold changes obtained from the limma-voom method of Law *et al.* (2014) implemented in the `limma` package of Ritchie *et al.* (2015)^1^. An example gene selection of 569 genes is highlighted in red. It corresponds to genes whose *p*-value is less than 10^-3^ and fold change larger than 0.5 in absolute value. The Adaptive Simes method ensures that with probability larger than 90%, the proportion of false discoveries (FDP) is less than 0.14. It also ensures that the FDP among the subset of genes with positive fold change is less than 0.14, and that the FDP among the subset of genes with negative fold change is less than 0.63. As already noted, the proposed bounds can be computed for multiple, arbitrary genes subsets without comprising their validity. Here again, the bounds provided by the ARI method are less tight than the Adaptive Simes method, as illustrated in Table 1.

## 6. Statistical performance for DE studies

### 6.1. Existing post hoc inference methods

The first post hoc inference methods introduced in were not adaptive to the dependence between tests, since they were obtained from probabilistic inequalities:

- The **Simes** bound was first proposed in Goeman and Solari (2011) together with a quadratic algorithm ((*O*(*m*^2^))). It has been implemented in the R package `cherry`
- A slightly sharper version of the Simes bound has been introduced in Goeman *et al.* (2019), together with an algorithm of linearithmic complexity. This method is known as **ARI** for “All resolution inference” and implemented in the R package `hommel`.

The idea of using randomization to obtain sharp risk control is not new in the multiple testing literature. In particular, resampling or permutations have been used to control the Family-Wise Error Rate (FWER) (Ge *et al.*, 2003; Westfall and Young, 1993) and the *k*-FWER (Romano and Wolf, 2007). For post hoc inference:

- The **Adaptive Simes** method described in this paper exploits sign-flipping and permutation-based approaches introduced in Blanchard *et al.* (2020, 2021) in order to build post hoc bounds. It is implemented since 2017 in the R package `sanssouci`.
- A closely related approach called **pARI** has recently been proposed by Andreella *et al.* (2020) for the analysis of neuroimaging data. It is implemented in the R package `pARI`.

Both pARI and Adaptive Simes rely on the calibration method described in Section 3, combined with the interpolation bound (3). However, two important differences between Adaptive Simes (R package `sanssouci` and pARI are worth to be mentioned. First, `sanssouci` implements the linearithmic time complexity algorithm descrined in Section 4. In contrast, the algorithm used to calculate the post hoc bound after calibration is the “naive” interpolation algorithm described in the beginning of Section 4, which has a quadratic complexity for a single set.

Secondly, contrary to pARI, `sanssouci` implements a step-down principle in order to adapt to the unknown quantity of signal (or, equivalently, to the proportion *π*_0_ of true null hypotheses). This makes the Adaptive Simes method slightly sharper than pARI. In the numerical results below, we have used instead of the instead of the `pARI` package, the single-step version of the Adaptive Simes method, which is mathematically equivalent, algorithmically faster, and implemented in the `sanssouci` package. The main features of existing post hoc bounds are summarized in Table 2.

**Table 2:**
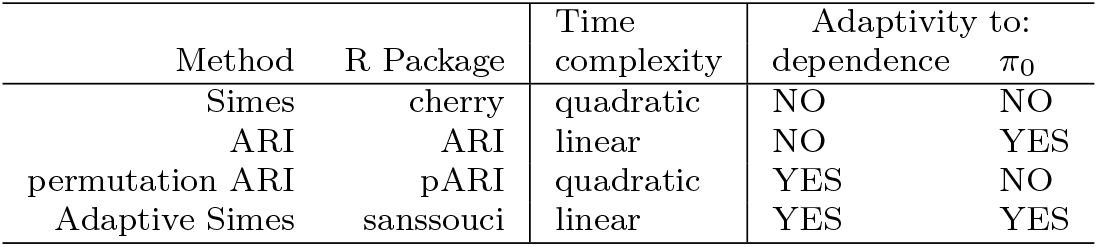
Main features of existing post hoc inference methods and software.

**Table 3.** *Post hoc bounds on BLCA data set for the four compared methods. Left panel: Gene selections from Figures 2 and 5, with target FDP set to* 0.1. *Right panel: gene selection for the volcano plot (Figure 3).*

|  | Confidence curve (Fig. 2) |  |  | Volcano plot (Fig 3) |  |  |
| --- | --- | --- | --- | --- | --- | --- |
|  | \|S\| | TP<sub>α</sub>(S) | FDP<sub>α</sub>(S) | \|S\| | TP<sub>α</sub>(S) | FDP<sub>α</sub>(S) |
| Adaptive Simes | 1042 | 958 | 0.1 | 569 | 492 | 0.135 |
| Adaptive Simes (single step) | 1042 | 938 | 0.1 | 569 | 490 | 0.139 |
| ARI | 781 | 703 | 0.1 | 569 | 456 | 0.199 |
| Simes | 757 | 682 | 0.1 | 569 | 452 | 0.206 |

### 6.2. Evaluation framework

The mathematical validity of the post hoc bounds considered in this paper has been proved in Blanchard *et al.* (2020), where their numerical performance has also been illustrated by experiments on synthetic data. The goal of this section is to complement these results by numerical experiments based on gene expression data, which are more realistic for the purpose of DE studies.

#### Data set generation

In the absence of a gold standard data set where one would know which genes are truly DE or not, we created such data sets as follows. Starting from a *n* × *m* gene expression data set *X*, where each row corresponds to a gene and each column to an experiment or statistical observation, we have

1. partitioned the observations into two groups of size *n*_0_ and *n*_1_, such that *n*_0_ + *n*_1_ = *n*;
2. partitioned the genes into *m*_0_ null genes and *m*_0_ non-null genes, with *m*_0_ + *m*_1_ = *m*
3. modified the expression of the non-null genes in group 1 by shifting or scaling the corresponding submatrix of *X* of size *n*_1_ × *m*_1_.

This process results in a perturbed gene expression data set *Y* where the null and non-null genes are known. Following Blanchard *et al.* (2020), we have quantified, for a set of such experiments, estimates of the risk (JER) and of the power of each method considered, for each value of the target risk *α*. The JER results are presented in Section 6.3. The power results, which are highly consistent with the JER results, are postponed to Supplementary Materials.

The empirical risk of a given method is estimated by the proportion of experiments for which the corresponding confidence curve on the false positives is now always below the actual number of false positives. This quantity is the empirical counterpart of the JER defined in (2), and can be compared to the target risk *α*: JER is empirically controlled if the empirical JER is lower than *α*, and the closer it is to *α*, the tighter JER control.

The parameters of such a numerical experiment are the proportion *π*_0_ = *m*_0_/*m* of null genes, and a measure of distance (or signal to noise ratio) between null and non-null genes. Section 6.3 reports the numerical results obtained for RNA sequencing data. We have also performed the same type of experiments with microarray data. The results are similar, and they are reported in Section D.4.

### 6.3. Results for bulk RNA sequencing data

Our starting point is the data set used in section 5. For this experiment we have selected only stage III samples, and performed the same filtering as in Section 5 on these samples only. We obtained a “null” data set (with no signal), consisting of 130 patients and *m* = 12, 418 genes, after applying the same process as described in Section 5 for filtering out unexpressed genes. The parameters of the experiments are set as follows. The proportion of null genes is set to *π*_0_ ∈ {0.8, 0.95, 1}. We have considered an multiplicative signal for differential expression: for each gene *g* among the *m*_1_ non-null genes, the original expression values of *g* are multiplied by a constant *s_g_* for *n*_1_ of the *n* observations, where *s_g_* is drawn uniformly between 1 and a signal to noise (SNR) parameter. The SNR value is set to 1 (no signal), 2 or 3 (weak to strong signal). We have used a two-sided Wilcoxon rank sum test for comparing the two groups.

The results are summarized by Figure 4, where the average empirical risk (JER) achieved across 1000 experiments is plotted (together with 95% confidence curves) against the target risk *α* for the methods described in Section 6.1. In particular, pARI is represented by the “Adaptive Simes (single step)” method.

**Fig 4.**
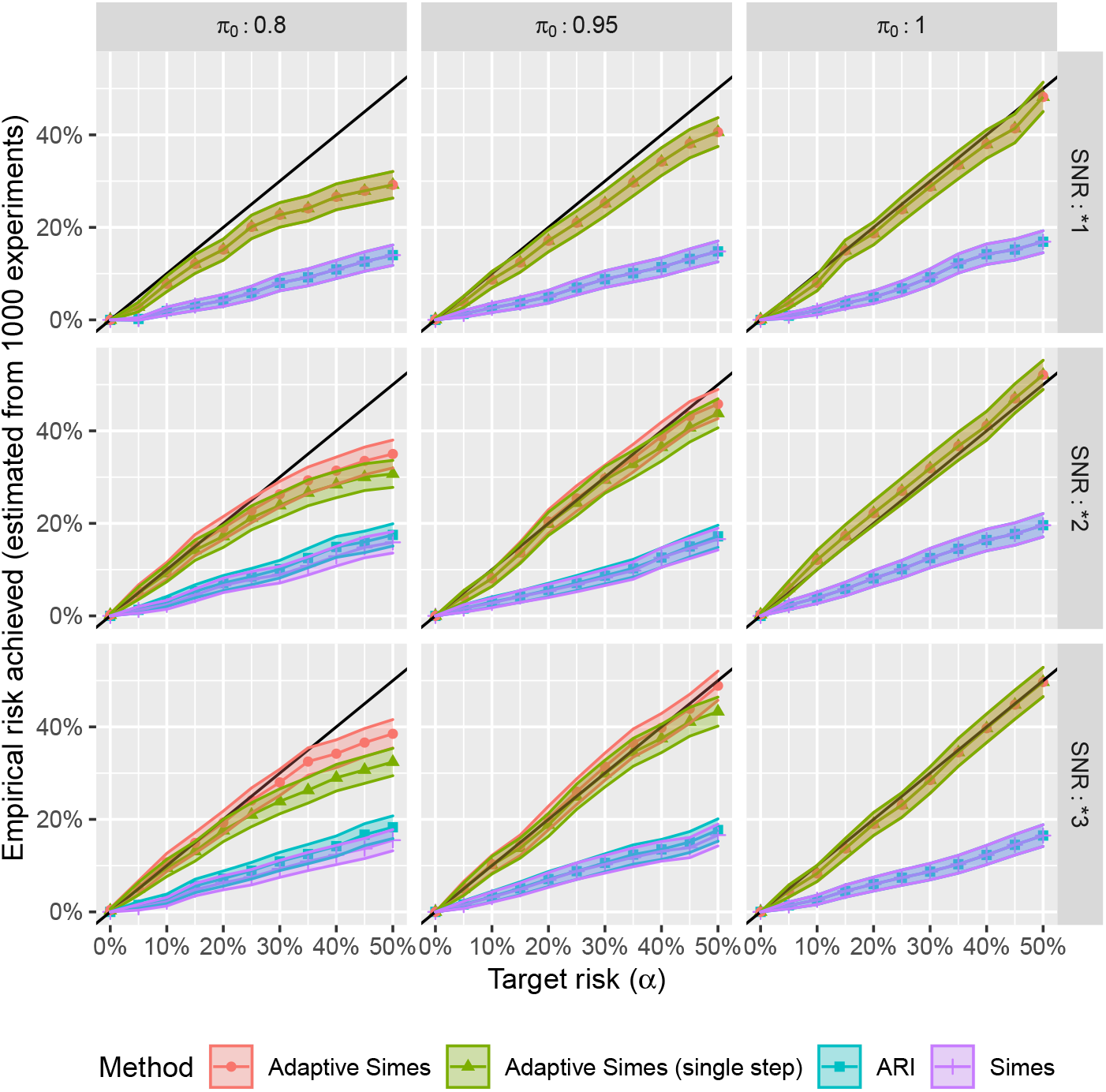
*Validity and compared tightness of the post hoc bounds on RNA-seq based numerical experiments. The average empirical JER achieved across* 1000 *experiments is plotted (together with* 95% *confidence curves) against the target risk* α *for all considered methods. Each panel corresponds to a combination of the parameters π*_0_ *and SNR.*

**Fig 5.**
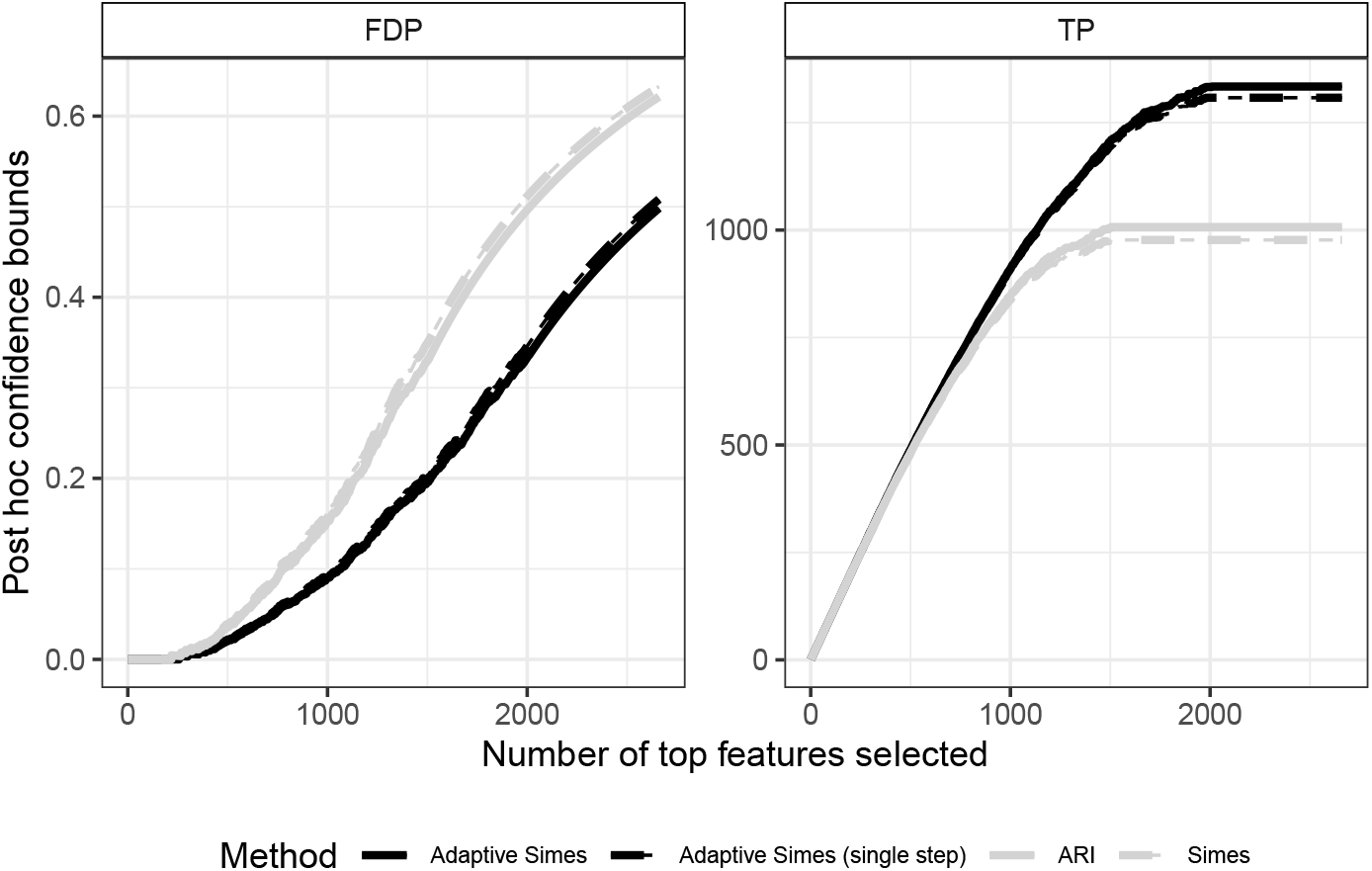
90% *confidence curves on “top k” lists for the Urothelial bladder carcinoma data set. Left: upper bound on the False Discovery Proportion (FDP); right: lower bound on the number of true positives (TP). Adaptive methods (black curves) outperforms non-adaptive ones (gray curves).*

Each panel corresponds to a combination of the parameters *π*_0_ ∈ {0.8, 0.95, 1} (in columns) and SNR ∈ {1, 2, 3} (in rows). The JER is controlled for all methods and all parameter combinations, since all curves are below the diagonal. The risk for the Adaptive Simes methods is substantially closer to the target risk than for the parametric Simes methods (Simes and ARI). This illustrates the systematic gain in tightness provided by the calibration method described in Section 3. We also note that the gain obtained from the adaptation to *π*_0_ is very small, and even negligible for *α* ≤ 0.2. Indeed, the Simes and ARI methods are essentially indistinguishable from each other, and the same holds for the single-step and step-down Adaptive Simes methods. These results are also confirmed by those of the power assessment, which are given in Supplementary Materials.

## 7. Discussion

This paper advocates for the use of post hoc inference in DE studies, which provide more interpretable statistical guarantees than classical inference based on the False Discovery Rate. The methods proposed in this paper make it possible to obtain post hoc bounds that are both fast to compute, and powerful (in the sense of the proportion of true signal recovered). The resulting improvement over the state-of-the-art is illustrated by realistic numerical experiments based on RNAseq and microarray data. These methods are implemented in the opensource R package `sanssouci`. The code used for the numerical experiments of this paper and to generate the figure is also provided, making the results of this paper reproducible and exploitable for future research.

The methods proposed in this paper and their implementation in the R package `sanssouci` are generic, in the sense that they can be used with reference families (or *templates*) of arbitrary shape. The most natural choice is that of the Simes family, as it is closely related both to FDR control and to the first post hoc bounds introduced by Goeman and Solari (2011). This results in the Adaptive Simes method, which has been used in the numerical experiments reported in this paper. An interesting perspective of this work is to compare the performance of other templates. Our experience in DE studies indicates that improving on the Simes family is challenging; similar conclusions have been reported in Andreella *et al.* (2020) for the analysis of fMRI data.

While this paper focuses on DE studies, these methods and our implementation are applicable to any practical situation involving multiple two-sample (or one-sample) tests. Such situations are frequent in genomics (differential expression, differential splicing, differential methylation) but also in neuroimaging, which is another field where post hoc inference methods have been introduced (Rosenblatt *et al.*, 2018). However, for studies have more complex designs such as multi-sample comparisons or studies including covariates, the calibration-based approach proposed here cannot be applied directly. Extensions of the present work to the problem of testing parameters of a general linear model is another interesting perspective.

## Funding

This work has been supported by ANR-16-CE40-0019 (SansSouci), by Fondation Catalyses at Université Paul Sabatier, and by the Mission for Transversal and Interdisciplinary Initiatives (MITI) at CNRS through the DDisc project.

## Appendix A: Calibration algorithm

The calibration algorithm described in Section 3 and illustrated in Figure 1 is formalized in Algorithm 2.

**Algorithm 2.**
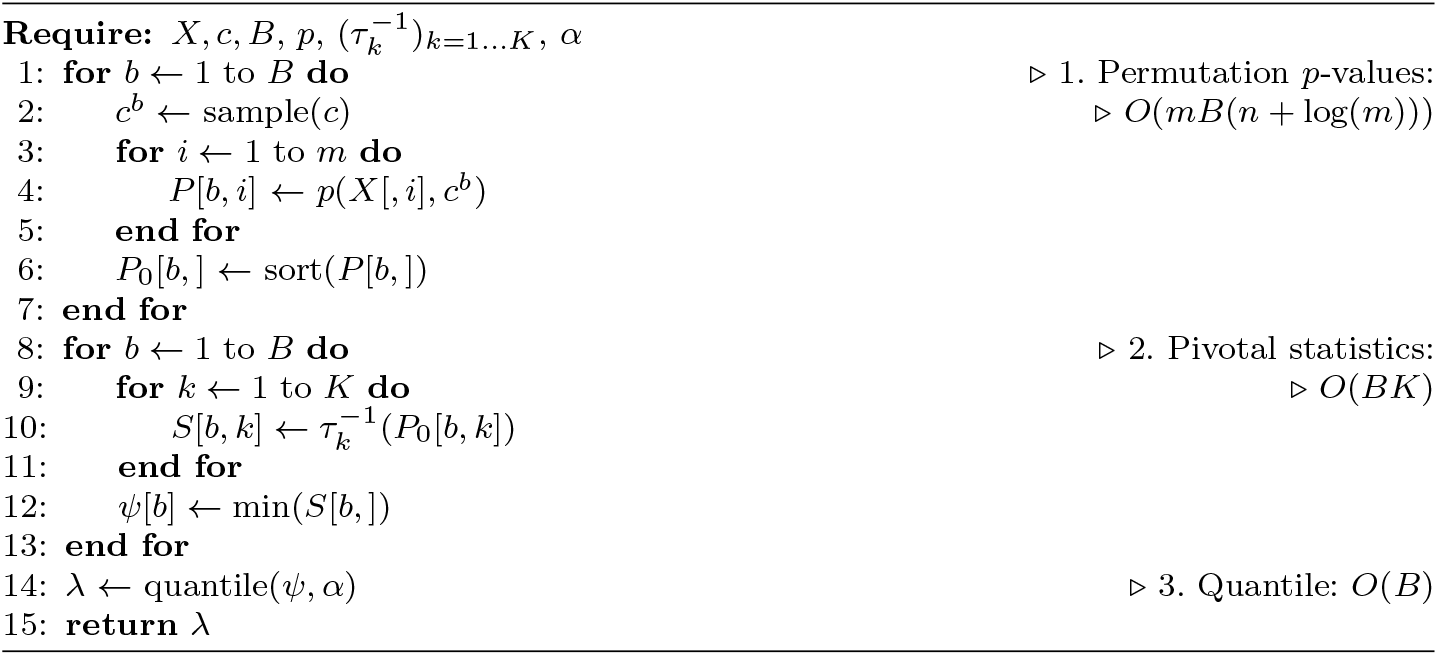
Calibration

## Appendix B: Proof of Proposition 1 (interpolation-based post hoc bound)

We denote by 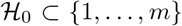 the (unknown) subset of true null hypotheses. Then for a given subset *S* of genes, the number of false positives in *S* can be written as 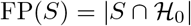.

#### Proof of Proposition 1.

The proof relies on the following simple observation: for any subsets *S* and *R* of {1,…, *m*}, we have

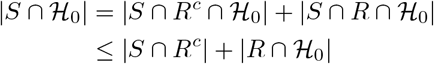

Letting *R_k_* = {*i* ∈ {1,∈,*m*}, *p_i_* ≤ *t_k_*} be the set of genes whose *p*-value is less than *t_k_* for 1 ≤ *k* ≤ *K*, we note that

i. 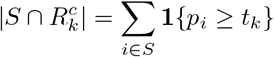
ii. 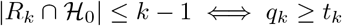

By (2), it holds with probability greater than (1–*α*) that for all 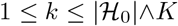. Therefore, there exists an event of probability greater than (1–*α*) such that for any *S* ⊂ {1,…,*m*},

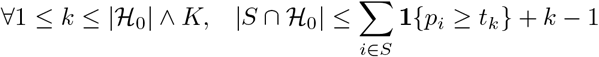

which concludes the proof.

## Appendix C: Validity of Algorithm 2 (linear time interpolation bound)

A first step is to rewrite the bound 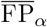 on arbitrary subsets of *S* as a minimum over |*S*| items:

#### Lemma 1.

Let *S* ⊂ {1,…,*m*}. Then for any *R* ⊂ *S*,

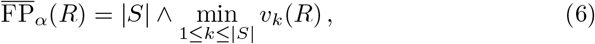

*where v_k_* (*R*) = ∑_*i*∈*R*_ **1**{*p_i_* ≥ *t*_*k*Λ*K*_} + *k* – 1.

*Proof of Lemma 1.* Let *R* ⊂ *S* ⊂ {1,…,*m*}. Let 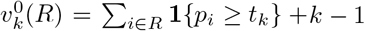 for *k* = 1,…,*K*. With this notation, we have

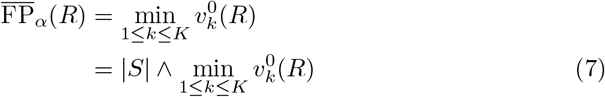

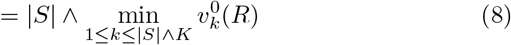

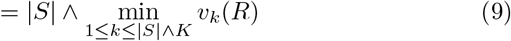

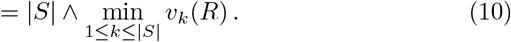

Above, Equation (7) holds since 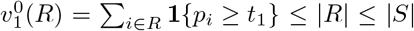. Equation (8) is obvious if |*S*| ≥ *K*; if |*S*| < *K* then for *k* > |*S*|∧*K*, 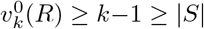. Equation (9) holds since 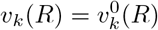 for *k* ≥ |*S*|∧ *K*(= *K*), we have *t*_*k*∧*K*_ = *t_K_* which implies that *v_k_*(*R*) = (*k* – *K*) + *v_k_*(*R*) ≤ *v_k_*(*R*).

We are now ready to prove Proposition 2.

### Proof of Proposition 2.

We consider the nested subsets *S_i_* for *i* = 1,…, *s*. By Lemma 1, we have

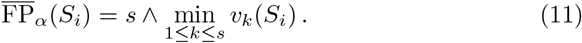

If *p_i_* ≥ *t*_*s*∧*K*_, then *κ_i_* = *s*, which implies that

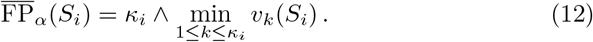

If *p_i_* ≥ *t*_*s*∧*K*_, then *κ_i_* < *s*. This implies that *v_k_*(*S_i_*) ≥ *κ_i_* for all *k* ∈ {*κ_i_* + 1,∈,*s*}, and *v*_*κ_i_*+1_(*S_i_*) = *κ_i_*. Therefore, we have

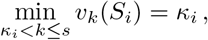

so that (12) holds as well. We conclude by noting that for *k* ≤ *κ_i_*,

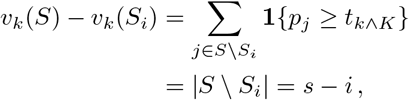

since for *j* ∈ *S* \ *S_i_*, *p_j_* ≥ *p_j_* ≥ *p_i_* > *t*_*κ_i_*∧*K*_ ≥ *t*_*k∧K*_.

## Appendix D: Additional results

### D.1. Comparison between existing post hoc bounds

This section complements the results presented in Section 5 of the main paper, by providing the results of the (non-adaptive) Simes method and the single-step adaptive Simes method (which corresponds to the pARI method). All methods are described in Section 6.1. This Figure illustrates the fact that for DE studies, the adaptation to dependency provided by the calibration method described in Section 3 (black vs gray curves) yields a more substantial improvement than the adaptation to the proportion of true null hypotheses (dashed vs solid curves).

### D.2. Comparison between limma-voom and Wilcoxon *p*-values

As explained in Section 3, the theoretical results underlying our calibration method require the test statistic (or *p*-value) for each gene to depend on the data only via the expression measurements associated with this gene. Hovever, the most commonly used statistical tests in DE studies with RNAseq data (DE-seq2 Love *et al.* (2014), edgeR Chen *et al.* (2014) and limma-voom Ritchie *et al.* (2015)) do not formally meet this assumption. In particular, these methods are using moderated variance estimators that borrow information from all genes, which is crucial when dealing with low sample sizes. Let us emphasize that, by the very nature of post hoc bounds and is illustrated in Figure 3, our methods can still be used to evaluate the number of false positives in any gene selection, *including selections obtained from the above-mentioned methods.*

For completeness, Figure 6 provides a graphical comparison between the *p*-values obtained by the limma-voom and the Wilcoxon rank sum test for the bladder urothelial carcinoma data set. This plot illustrates the consistency between these *p*-values.

**Fig 6.**
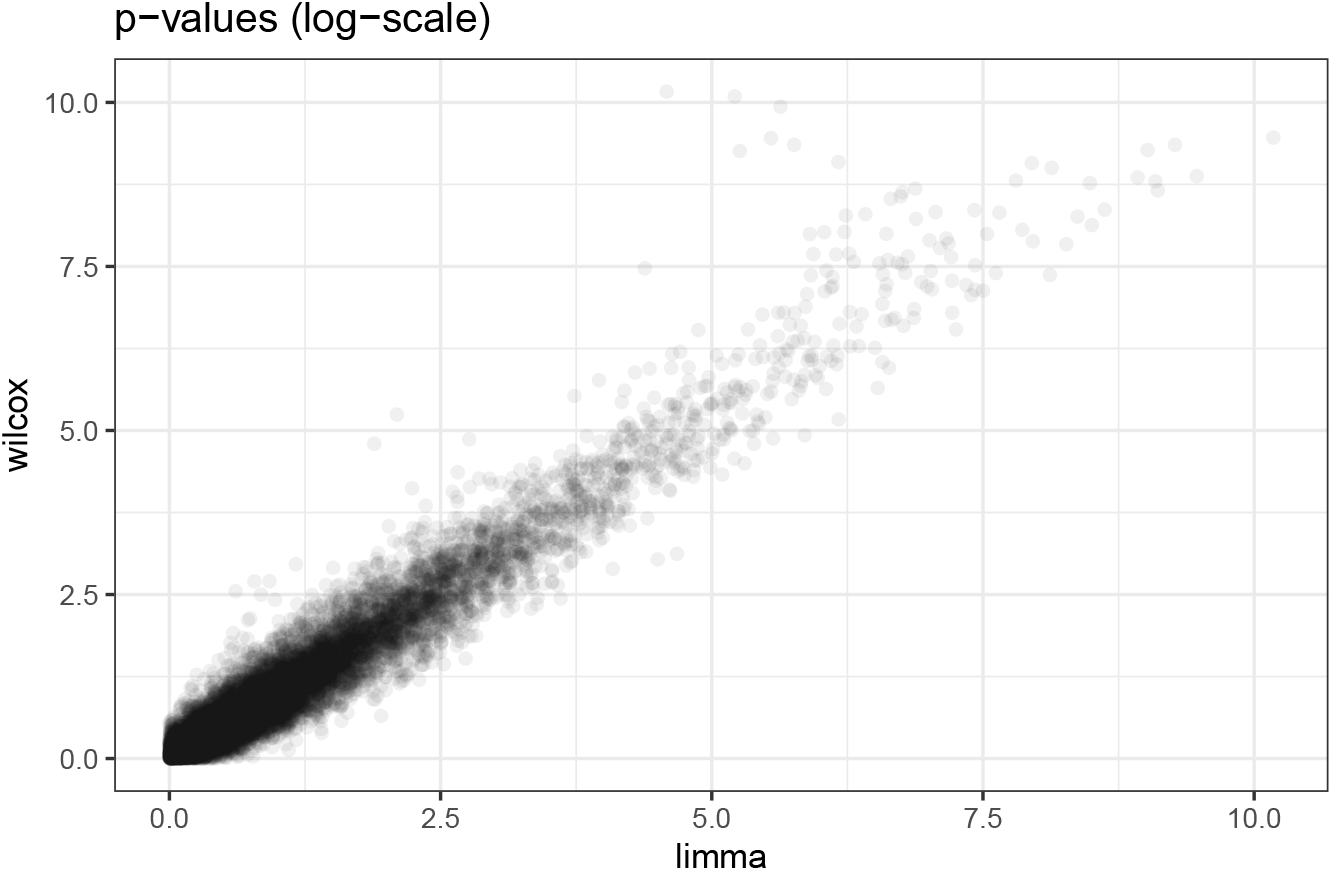
Comparison between limma-voom and Wilcoxon *p*-values on the BLCA data set.

### D.3. Power assessment for RNAseq data

For a given subset *S* of genes, the power of a method providing the post hoc bound 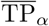 is defined in Blanchard *et al.* (2020) as

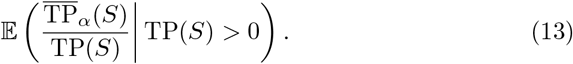

This quantity corresponds to the expected proportion of signal in *S* actually recovered by the method considered. Again, it is estimated by its empirical counterpart, that is, the average proportion of signal in *S* recovered over those experiments for which some signal was actually present in *S*. We considered four different gene selections *S* for estimating the power as defined in (13):

**BH_**05: the set genes selected by the Benjamini-Hochberg procedure Benjamini and Hochberg (1995) at level 5%
**first_100: the 100 genes with lower *p*-value**
**p_**05: the genes whose *p*-value is less than 5%
**H:** all genes in the data set.

Figure 7 displays the empircal power for the following choice of parameters: *π*_0_ = 0.8 and *SNR* ∈ {1.5, 3} The power of the Adaptive Simes methods is higher than the one of parametric Simes methods. This is coherent with the increased tightness of the Adaptive Simes bounds relative to their parametric counterpart already observed in Figure 4. We also note than the adaptivity to *π*_0_ does not make a difference in terms of power: the Simes and ARI methods are essentially indistinguishable from each other, and the same holds for the single-step and step-down Adaptive Simes methods.

**Fig 7.**
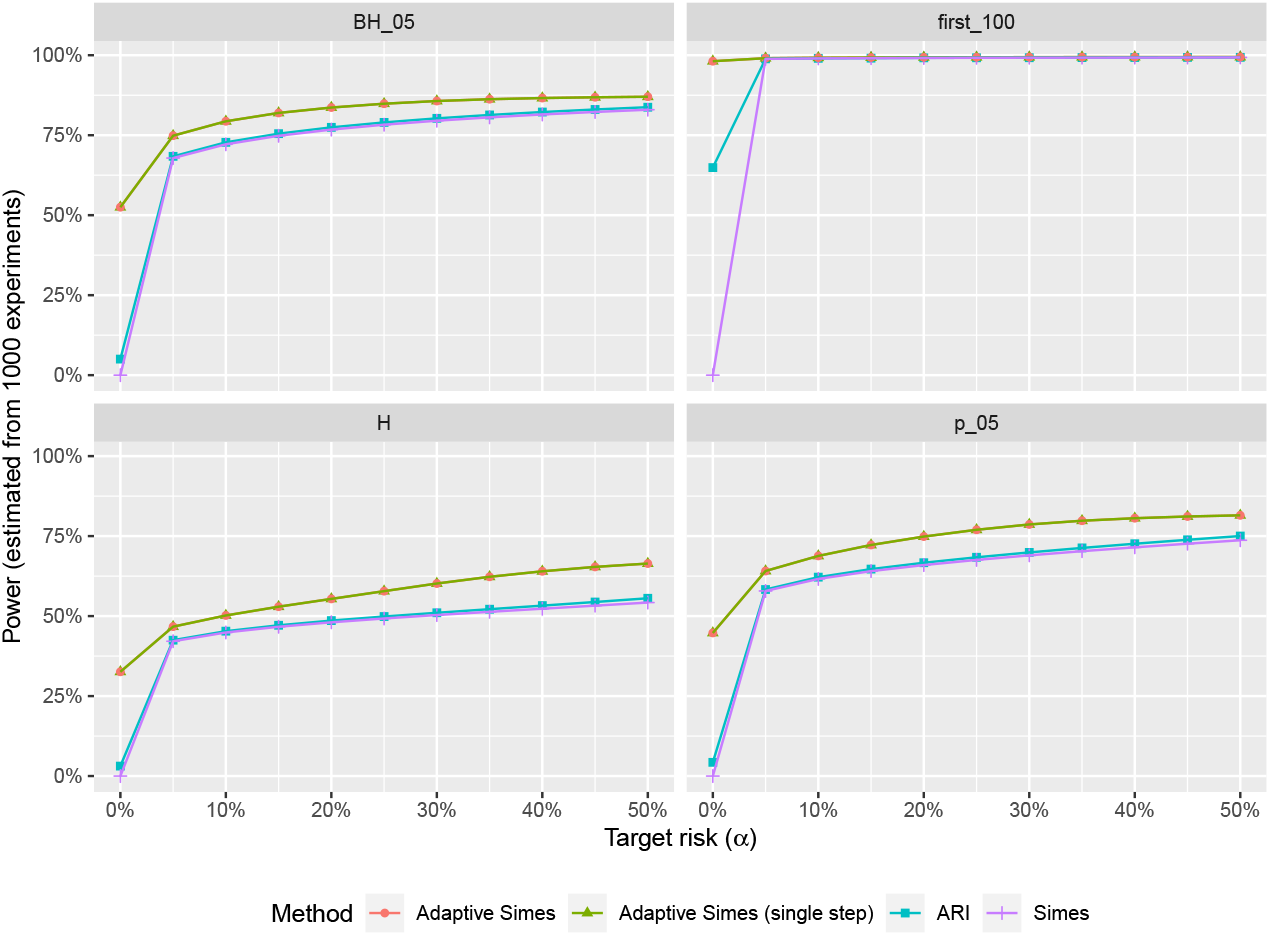
*Empirical power of four methods, as a function of the target risk (JER) level. Simes-based methods are outperformed by their adaptive couterpart. Each panel corresponds to a different gene selection (as described in main text).*

### D.4. Statistical valididy: results for microarray data

We considered the GSE19188 Hou *et al.* (2010) data set available from GEO Barrett *et al.* (2012) and the R package GSEABenchmarker Geistlinger *et al.* (2020). This data set consists of 91 non-small cell lung cancer tissue samples and 62 normal samples. Each of these observations corresponds to a vector of *m* = 21, 407 gene expression values.

The parameters of the experiments have been set as follows. The proportion of null genes is set to *π*_0_ ∈ {0.5, 0.8, 1}. We have considered an additive signal for differntial expression: the expression level for the *m*_1_ non-null genes are then shifted by a constant value for *n*_1_ of the *n* observations. We have used a twosided Welch test for comparing the two groups. The results are summarized by Figure 8.

**Fig 8.**
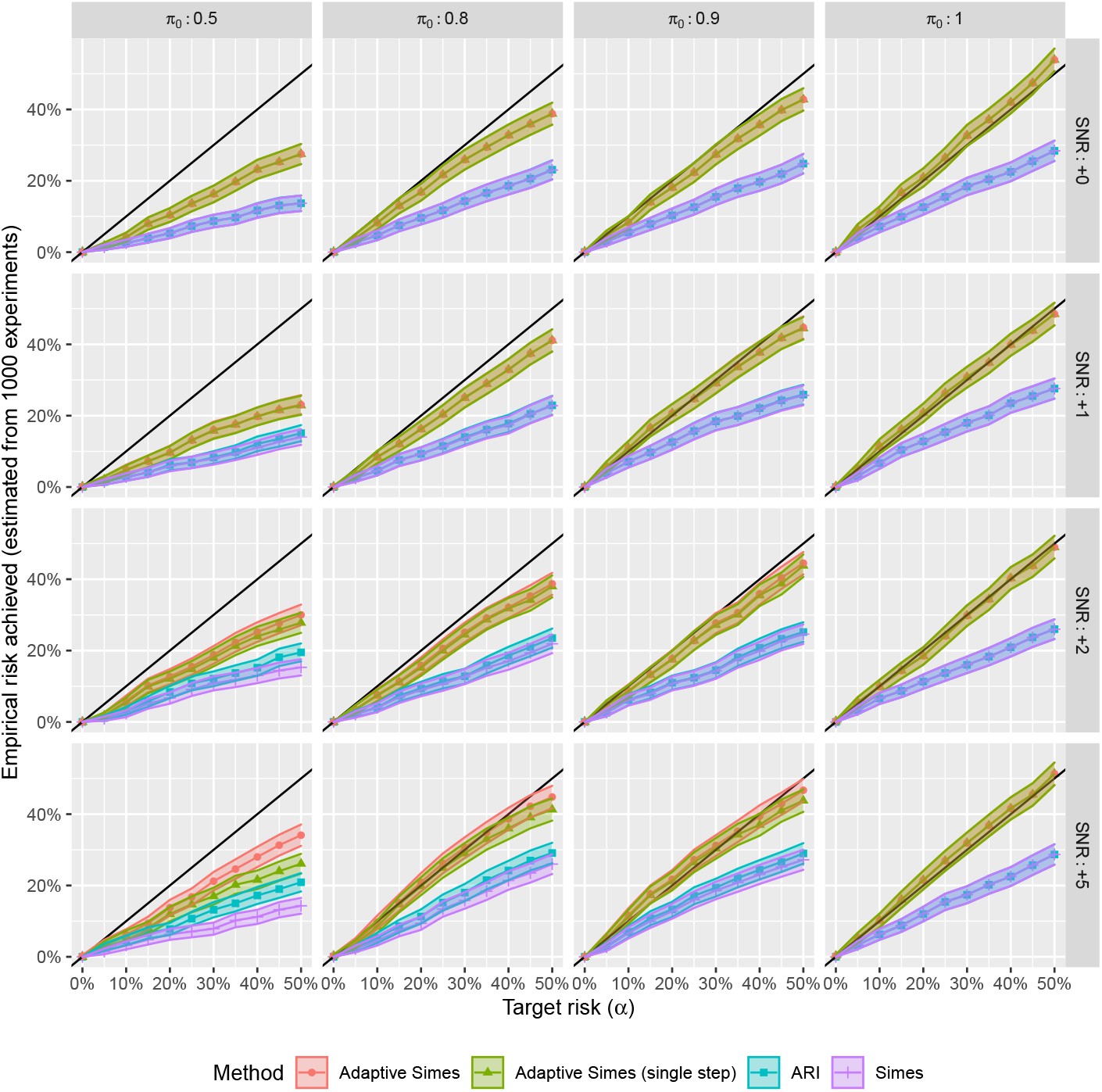
*Validity and compared tightness of the post hoc bounds on microarray-based numerical experiments. The average empirical JER achieved across *1000* experiments is plotted (together with *95%* confidence curves) against the target risk *α* for all considered methods. Each panel corresponds to a combination of the parameters π*_0_ *and SNR*.

The conclusions are identical to those obtained for RNAseq-based experiments in the main paper: JER is controlled for all methods and all parameter combinations, and the risk for the Adaptive Simes methods is substantially closer to the target risk than for the parametric Simes methods (Simes and ARI), illustrating a substantial gain provided by the calibration method described in Section 3. The gain obtained from the adaptation to *π*_0_ is negligible, except for case with strong and dense signal (*π*_0_ = 0.5 and SNR= 5), which is the less realistic scenario for DE studies.

## Appendix E: Implementation notes

The calibration method described in Section 3 has been available since 2017 within the R package `sanssouci`. Algorithm 1 has been available since 2016 within `sanssouci`^2^. The original implementation of Algorithm 1 had *O*(*K* ∨ *s*) time and space complexity, i.e. *O*(*m*) worse case complexity. It has been improved to *O*(*s*) complexity in 2021 by adding the lines 1-8.

## Footnotes

1 A quantitative comparison of the Wilcoxon and limma-voom *p*-value is provided in Figure 6. It illustrates the coherence of the two methods for identifying DE genes.

2 See https://github.com/pneuvial/sanssouci/commit/a83082b.

## Notes

### Competing Interest Statement

The authors have declared no competing interest.

